# LeafyVGG-16: Transfer Learning for Plant Disease Detection with Cyber Risk Analysis

**DOI:** 10.64898/2026.05.13.724946

**Authors:** Natwange Chiwele, Eddie Sweeney, Kabir Hossain

## Abstract

Plant disease detection using deep learning is essential for precision agriculture, enabling early and automated crop health monitoring. This study proposes an end-to-end transfer learning pipeline, LeafyVGG-16, for multi-class classification of plant diseases and nutrient deficiencies using a tomato leaf dataset. The framework integrates data preprocessing, augmentation, and a VGG-16 backbone with a two-stage fine-tuning strategy. The proposed model is evaluated against CNN, DenseNet-121, Inception-V3, EfficientNetB0, and ResNet-50, achieving an accuracy of 0.93 with precision, recall, and F1-scores of 0.93, 0.90, and 0.92, respectively. These results demonstrate the effectiveness of transfer learning for fine-grained plant disease recognition. We further evaluate model robustness under adversarial cyber attacks to assess deployment reliability in agricultural systems. Under Fast Gradient Sign Method (FGSM) attacks (*ϵ* = 0.01– 0.05), the model shows an accuracy drop of 1%–7.5%, while Projected Gradient Descent (PGD) attacks (*ϵ* = 0.05, step size = 0.005, 10 iterations) produce similar degradation, highlighting the model’s vulnerability to adversarial perturbations. These findings highlight potential security and reliability risks in AI-based agricultural decision-making systems. Future work will focus on improving robustness and cyber-resilience and extending this framework to other crops for secure and context-aware deployment in resource-constrained environments.

## I. Introduction

Ensuring sustainable agricultural production and global food security has become increasingly important due to rapid population growth, climate change, and increasing pressure on agricultural systems [1]–[3]. Plant diseases and nutrient deficiencies are major threats to global agricultural productivity, affecting crop yield, food quality, and economic stability [4]. Early and accurate identification of these conditions is essential for precision agriculture and sustainable farming [5]. Traditionally, plant disease diagnosis relies on manual inspection by experts, which is time-consuming, subjective, and difficult to scale for large agricultural systems [6].

Recent advances in deep learning and computer vision have made it possible to automatically detect plant diseases using convolutional neural networks (CNNs) [7]. Transfer learning methods [8], using pretrained models such as EfficientNetB0 [9], ResNet-50 [10], and InceptionV3 [11], have improved performance by reusing knowledge from large image datasets and reducing the need for large training data. However, plant disease recognition is still challenging because different diseases can look similar, the same disease can appear differently under different conditions, and real datasets are often imbalanced.

In this study, we propose a transfer learning-based method called LeafyVGG-16 for classifying plant diseases and nutrient deficiencies using the Tomato-Village Variant-a dataset. The system includes image pre-processing and data augmentation, followed by a VGG-16 model [12] pretrained on ImageNet [13], and a simple classification head adapted for plant disease recognition. A two-stage fine-tuning strategy is used to improve learning stability and performance. The overall proposed framework is illustrated in Fig. 1.

**Fig. 1.**
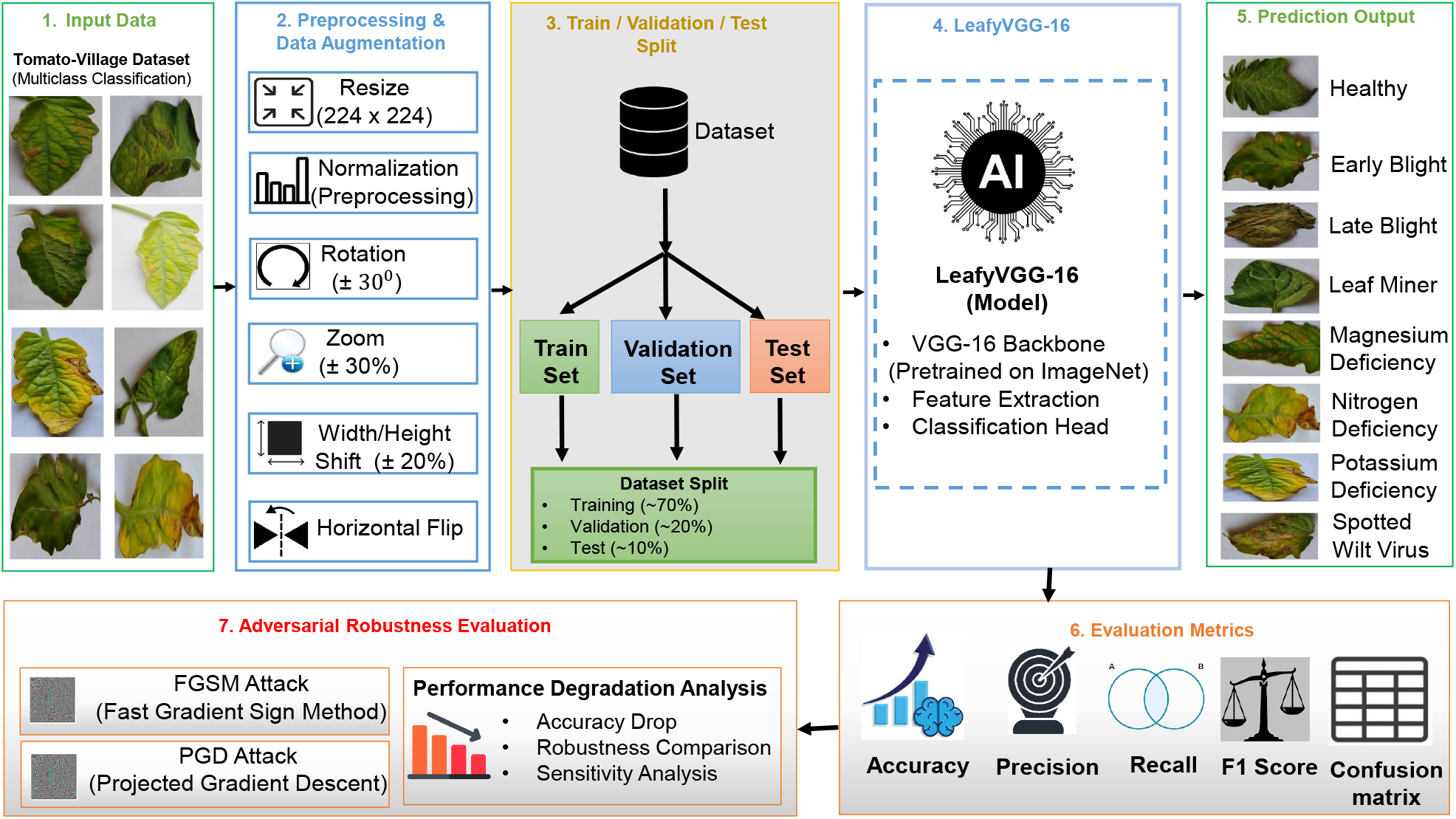
Overall framework of the proposed LeafyVGG-16 system for tomato leaf disease and nutrient deficiency classification. The pipeline includes image preprocessing and augmentation, dataset partitioning, transfer learning using a pretrained VGG-16 backbone, performance evaluation, and adversarial robustness assessment using FGSM and PGD attacks.

The proposed LeafyVGG-16 model is compared with several existing models, including a basic CNN, DenseNet-121 [15], InceptionV3, ResNet-50, and EfficientNetB0, using the same dataset and training setup. To ensure a fair comparison, all models are trained and evaluated under identical conditions.

Although deep learning models perform well under normal conditions, they are vulnerable to small, carefully crafted input changes known as adversarial perturbations. These changes are often indistinguishable to the human eye but can significantly affect predictions. In agricultural applications such as UAV-based monitoring and automated disease detection, this can lead to incorrect decisions, including missed disease detection or unnecessary pesticide use.

To analyze this issue, we evaluate the proposed LeafyVGG-16 model using two standard adversarial attacks: Fast Gradient Sign Method (FGSM) [16] and Projected Gradient Descent (PGD) [17]. FGSM generates single-step perturbations, while PGD applies iterative updates to create stronger attacks. This evaluation highlights the security and reliability of the model under adversarial conditions and demonstrates the need for more robust and cyber-secure plant disease detection systems.

The main contributions of this work are as follows:

- Development of an end-to-end transfer learning pipeline (LeafyVGG-16) for plant disease classification.
- Fine-tuning of a VGG-16 backbone with a custom classification head using batch normalization and dropout.
- Implementation of state-of-the-art models from the literature and comparison with the proposed model.
- Evaluation of adversarial robustness using FGSM and PGD, and its impact on risk in real-world AI agricultural systems.

The remainder of this paper is organized as follows: Section I presents the introduction. Section II reviews related work on plant disease classification and transfer learning approaches. Section III describes the proposed methodology, while Section IV presents the experimental results, including the performance of the proposed LeafyVGG-16 model, comparative analysis, and robustness evaluation under adversarial attacks. Finally, Section V concludes the paper and outlines future research directions focusing on the development of cyber-resilient models for precision agriculture.

## II. Related Work

Deep learning has been widely used for plant disease detection because it can automatically learn features from images without manual feature engineering. Early methods relied on handcrafted features combined with traditional machine learning models, but these approaches struggled with variations in lighting, texture, and disease appearance [19].

With the development of convolutional neural networks (CNNs) [20], significant improvements have been achieved in plant disease classification. CNN-based models can learn spatial patterns directly from images and generally outperform traditional methods. However, training deep CNNs from scratch requires large datasets and high computational resources, which are often not available in agricultural applications.

To address this limitation, transfer learning has become a popular approach. Pretrained models such as ResNet-50, DenseNet-121, InceptionV3, and EfficientNetB0, trained on large-scale datasets like ImageNet, have been successfully adapted for plant disease recognition tasks. VGG-based architectures are also widely used due to their simple yet effective design for extracting fine-grained leaf features. These models have shown strong performance in classification tasks under controlled experimental settings.

Despite their success, most existing studies mainly focus on improving classification accuracy under clean and ideal conditions. Recent research has also explored more advanced architectures, including hybrid CNN and attention-based models, to improve feature representation and classification performance.

More recently, increasing attention has been given to the robustness and reliability of deep learning models. Studies in adversarial machine learning have shown that CNN-based models are vulnerable to carefully designed perturbations that can significantly degrade performance. Methods such as FGSM and PGD are commonly used to evaluate these vulnerabilities. However, in plant disease detection, only limited studies have systematically analyzed adversarial robustness, especially on real-world agricultural datasets.

Unlike existing work that primarily focuses on classification accuracy, this study integrates transfer learning-based plant disease classification with a systematic adversarial robustness evaluation using FGSM and PGD. This provides a more complete assessment of model reliability and highlights its relevance for cyber-resilient agricultural AI systems.

## III. Methodology

### A. Dataset Description

This study uses the publicly available Tomato-Village dataset [18], which is designed for real-world tomato disease detection in agricultural environments. Unlike controlled laboratory datasets such as PlantVillage [23], which are collected under ideal conditions, the Tomato-Village dataset contains images captured in natural field environments, making it more suitable for real-world agricultural deployment scenarios [24].

In this work, we use only the Variant-a subset, which is formulated for multi-class classification. The dataset includes eight classes: Early Blight, Late Blight, Healthy, Leaf Miner, Magnesium Deficiency, Nitrogen Deficiency, Potassium Deficiency, and Spotted Wilt Virus.

The dataset contains a total of 4,525 images, split into training (3,162 images, 69.88%), validation (902 images, 19.93%), and testing (461 images, 10.19%). The dataset exhibits noticeable class imbalance, with some disease and deficiency classes having significantly more samples than others, which reflects real-world agricultural conditions and increases classification complexity. The overall dataset split distribution is illustrated in Fig. 3, while the class-wise distribution is shown in Fig. 2.

**Fig. 2.**
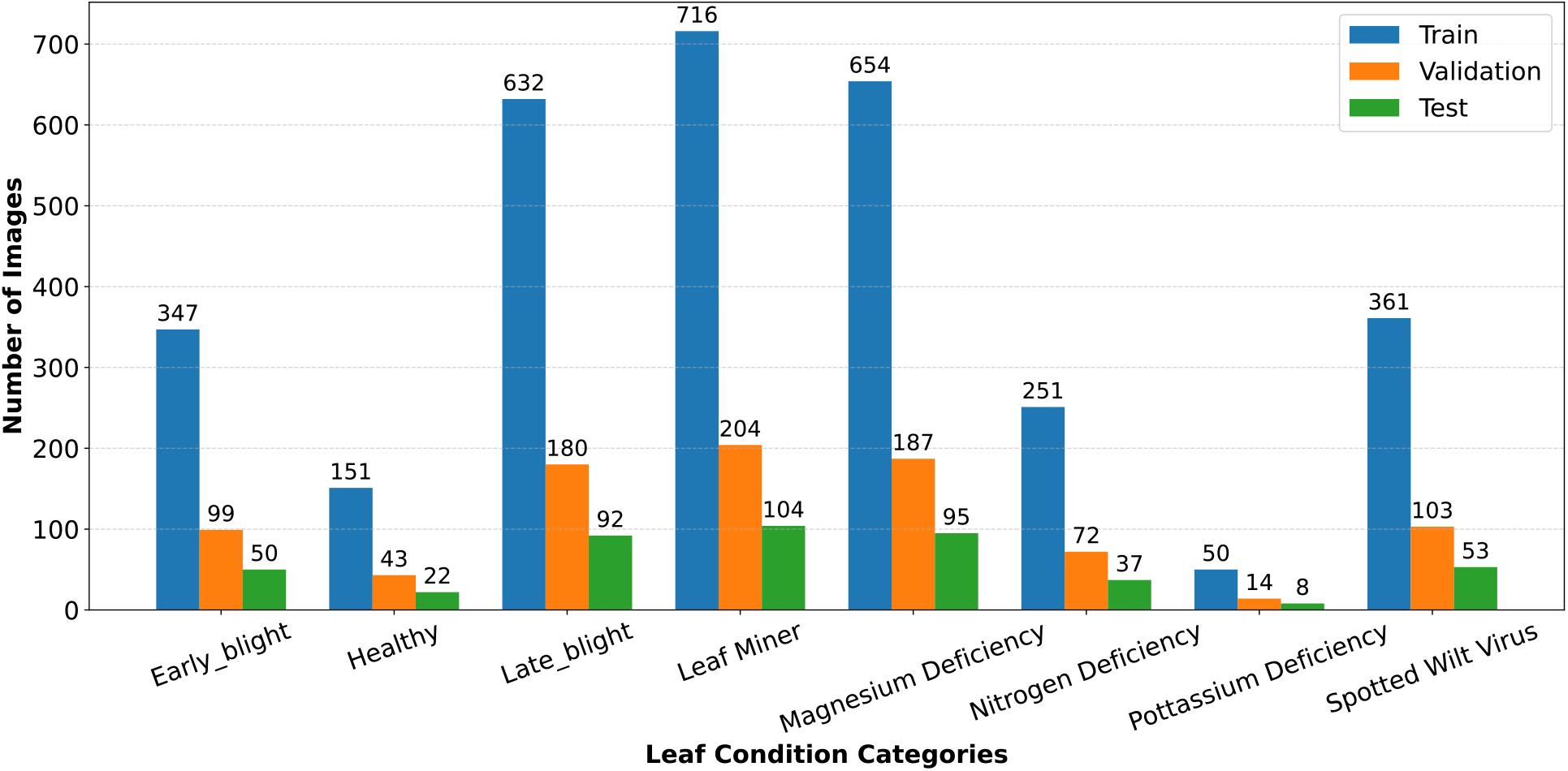
Class-wise distribution of the Tomato-Village Variant-a dataset across training, validation, and test sets.

**Fig. 3.**
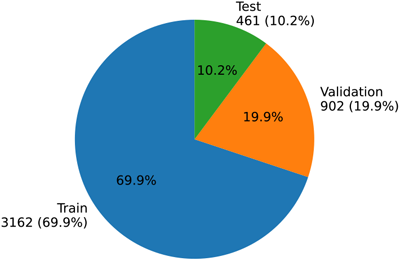
Training, validation, and test split distribution of the Tomato-Village Variant-a dataset.

To improve model generalization and reduce overfitting, data augmentation techniques such as random rotation and flipping [25], as well as scaling and zooming [26], are applied only to the training set. The validation and test sets remain unchanged to ensure unbiased evaluation. All images are resized to a fixed input resolution before being fed into the models.

### B. Proposed Model: LeafyVGG-16

The proposed LeafyVGG-16 framework is based on a transfer learning approach using the VGG-16 architecture as a feature extractor. The model leverages pretrained weights from ImageNet to capture general visual features and adapts them for plant disease and nutrient deficiency classification.

The extracted feature maps from the VGG-16 backbone are passed to a lightweight classification head. This head consists of a Global Average Pooling layer followed by fully connected layers with batch normalization and dropout for regularization, and a final Softmax layer for multi-class prediction. This design helps improve generalization while reducing overfitting.

A two-stage fine-tuning strategy is applied to improve performance. In the first stage, the pretrained backbone is frozen and only the classification head is trained. In the second stage, selected deeper layers of the backbone are unfrozen and fine-tuned with a lower learning rate to better adapt to the plant disease dataset. The overall architecture of the proposed LeafyVGG-16 framework is shown in Fig. 4.

**Fig. 4.**
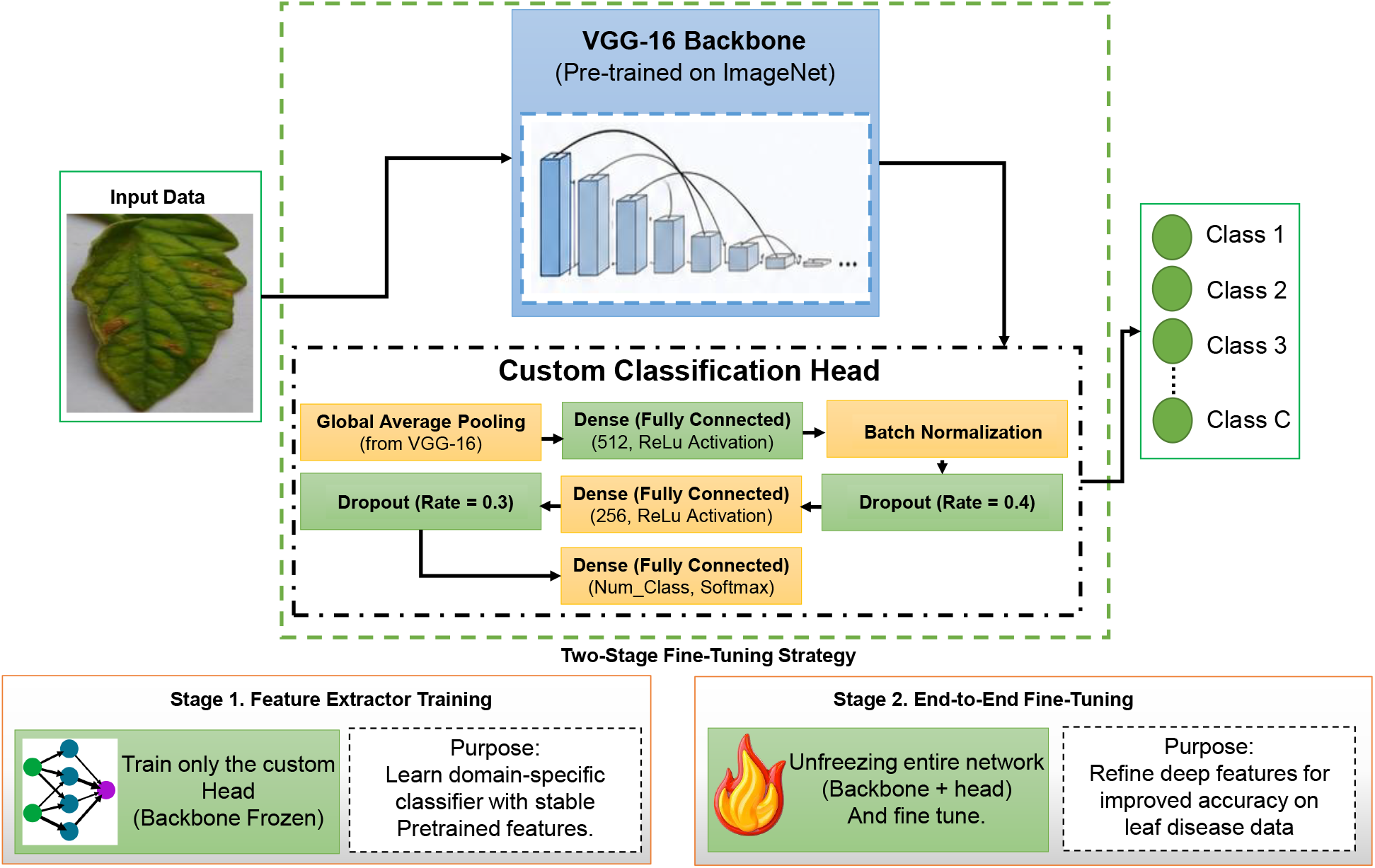
Proposed LeafyVGG-16 Framework

### C. Baseline Models for Comparison

The proposed LeafyVGG-16 framework is evaluated against several widely used deep learning architectures. These include pretrained transfer learning models, ResNet-50, DenseNet121, EfficientNetB0, and InceptionV3—along with a custom CNN trained from scratch. The pretrained models are adapted from ImageNet and fine-tuned for the plant disease classification task, while the custom CNN serves as a lightweight baseline to assess the effectiveness of transfer learning.

All models are trained and evaluated under identical experimental settings, including the same dataset split, input resolution, and training configuration, to ensure a fair comparison. Performance is reported using accuracy, precision, recall, and F1-score.

### D. Training Configuration

To ensure a fair comparison, all models including the proposed LeafyVGG-16, DenseNet-121, EfficientNetB0, InceptionV3, ResNet-50, and the custom CNN were trained under identical experimental settings. All experiments were conducted using TensorFlow with a fixed input image size of 224 × 224 pixels and a batch size of 32. The same training, validation, and test splits were used for all models to ensure consistent evaluation.

A two-stage training strategy was applied for all pretrained models. In the first stage, the pretrained backbone was frozen and only the classification head was trained using a learning rate of 1 × 10^−3^ for 50 epochs. In the second stage, selected layers of the backbone were unfrozen and fine-tuned with a reduced learning rate of 1 × 10^−5^ for up to 200 epochs.

All pretrained models were optimized using the Adam optimizer with categorical cross-entropy loss. To improve training stability, early stopping, learning rate reduction on plateau, and model checkpointing were used with standard configurations. Early stopping was based on validation loss with a patience of 70 epochs, and the learning rate was reduced by a factor of 0.5 when performance stopped improving.

In contrast, the custom CNN model was trained from scratch using a single-stage training strategy for 250 epochs using the same optimizer, loss function, and data augmentation strategy.

### E. Adversarial Robustness Evaluation

To evaluate the robustness of the proposed LeafyVGG-16 framework under adversarial conditions, we perform a systematic robustness analysis using two widely adopted gradient-based attack methods: the Fast Gradient Sign Method (FGSM) and Projected Gradient Descent (PGD). These attacks are applied to the trained model after standard training using the clean test dataset.

Let *x* denote the input image and *y* the corresponding ground-truth label. Adversarial examples are generated by introducing small perturbations to the input in the direction of the gradient of the loss function with respect to the input. FGSM generates adversarial samples using a single-step perturbation defined as:

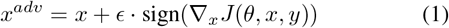

where *ϵ* controls the magnitude of the perturbation, *J*(*θ, x, y*) is the loss function, and ∇_*x*_ represents the gradient with respect to the input image.

In contrast, PGD is an iterative extension of FGSM, where adversarial samples are updated iteratively while constraining the perturbation within an *ϵ*-ball around the original input:

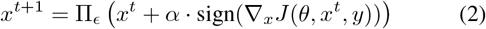

where *α* is the step size and Π_*ϵ*_ denotes projection onto the *ϵ*-bounded constraint space. The mathematical formulations for FGSM and PGD are adopted from the adversarial optimization framework introduced in [27].

## IV. RESULTS AND DISCUSSION

### A. Performance of the Proposed LeafyVGG-16 Model

The performance of the proposed LeafyVGG-16 model was evaluated using multiple quantitative metrics, including accuracy, precision, recall, F1-score, and confusion matrix. Experimental results demonstrate that the proposed framework achieved strong classification performance across the tomato leaf disease and nutrient deficiency categories while maintaining stable convergence during training.

The proposed LeafyVGG-16 model achieved an overall test accuracy and precision of 0.93, with a recall of 0.90 and F1-score of 0.92. These results indicate that the model effectively learned discriminative feature representations for identifying different disease symptoms and nutrient deficiencies from tomato leaf images.

Table I presents the detailed class-wise classification performance of the proposed model. High classification performance was achieved for most categories, particularly Magnesium Deficiency, Nitrogen Deficiency, and Late Blight, which obtained F1-scores of 0.97, 0.96, and 0.96, respectively. Similarly, the model achieved strong performance for Early Blight and Leaf Miner classes with F1-scores above 0.90. Comparatively lower performance was observed for the Healthy and Spotted Wilt Virus classes, mainly due to visual similarities between certain disease symptoms and healthy leaf patterns.

**TABLE I.** Class-wise performance of the proposed LeafyVGG-16 model.

| Class | Precision | Recall | F1-Score | Support |
| --- | --- | --- | --- | --- |
| Early Blight | 0.96 | 0.92 | 0.94 | 50 |
| Healthy | 0.89 | 0.73 | 0.80 | 22 |
| Late Blight | 0.95 | 0.97 | 0.96 | 92 |
| Leaf Miner | 0.88 | 0.94 | 0.91 | 104 |
| Magnesium Deficiency | 0.98 | 0.97 | 0.97 | 95 |
| Nitrogen Deficiency | 0.93 | 1.00 | 0.96 | 37 |
| Potassium Deficiency | 1.00 | 0.88 | 0.93 | 8 |
| Spotted Wilt Virus | 0.90 | 0.81 | 0.85 | 53 |
| **Overall Accuracy** |  |  | 0.93 |  |
| **Precision** |  |  | 0.93 |  |
| **Recall** |  |  | 0.90 |  |
| **F1-Score** |  |  | 0.92 |  |

The confusion matrix illustrated in Fig. 6 further demonstrates the classification capability of the proposed model. Most samples were correctly classified with limited inter-class confusion. Minor misclassifications were observed among visually similar categories, particularly between Healthy, Leaf Miner, and Spotted Wilt Virus classes.

**Fig. 5.**
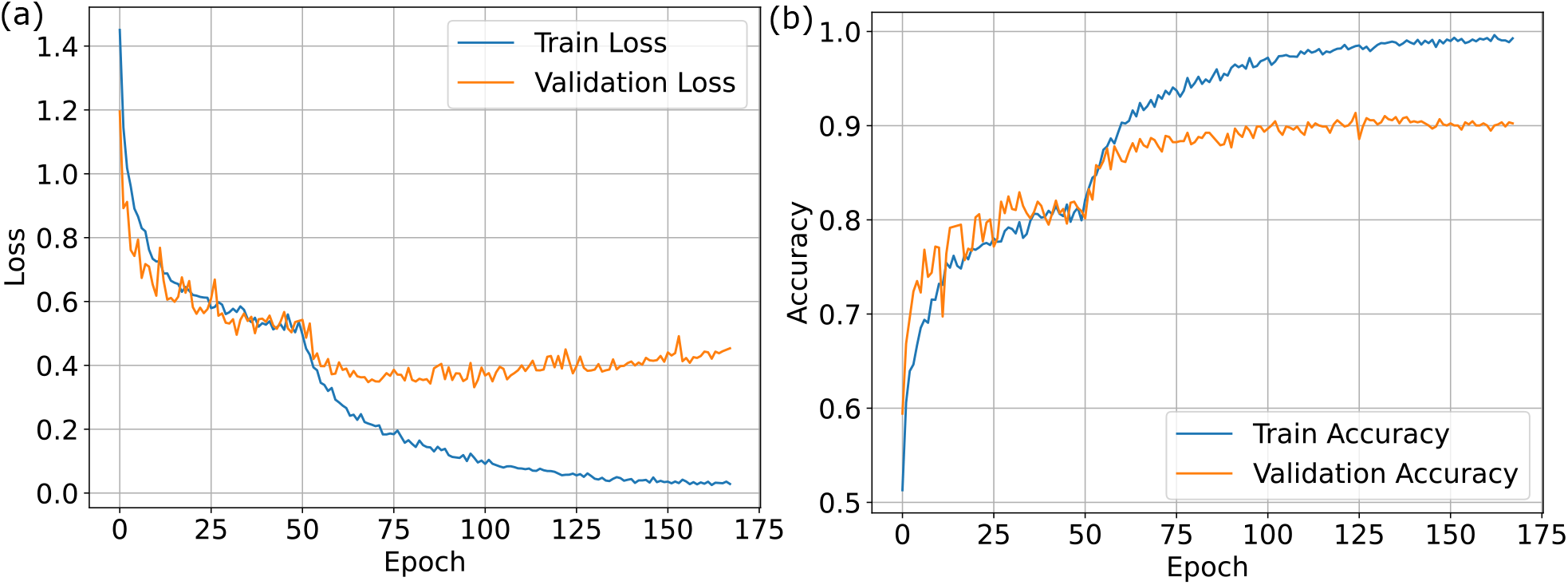
Training and validation performance curves of the proposed LeafyVGG-16 model: (a) accuracy curve and (b) loss curve across training epochs.

**Fig. 6.**
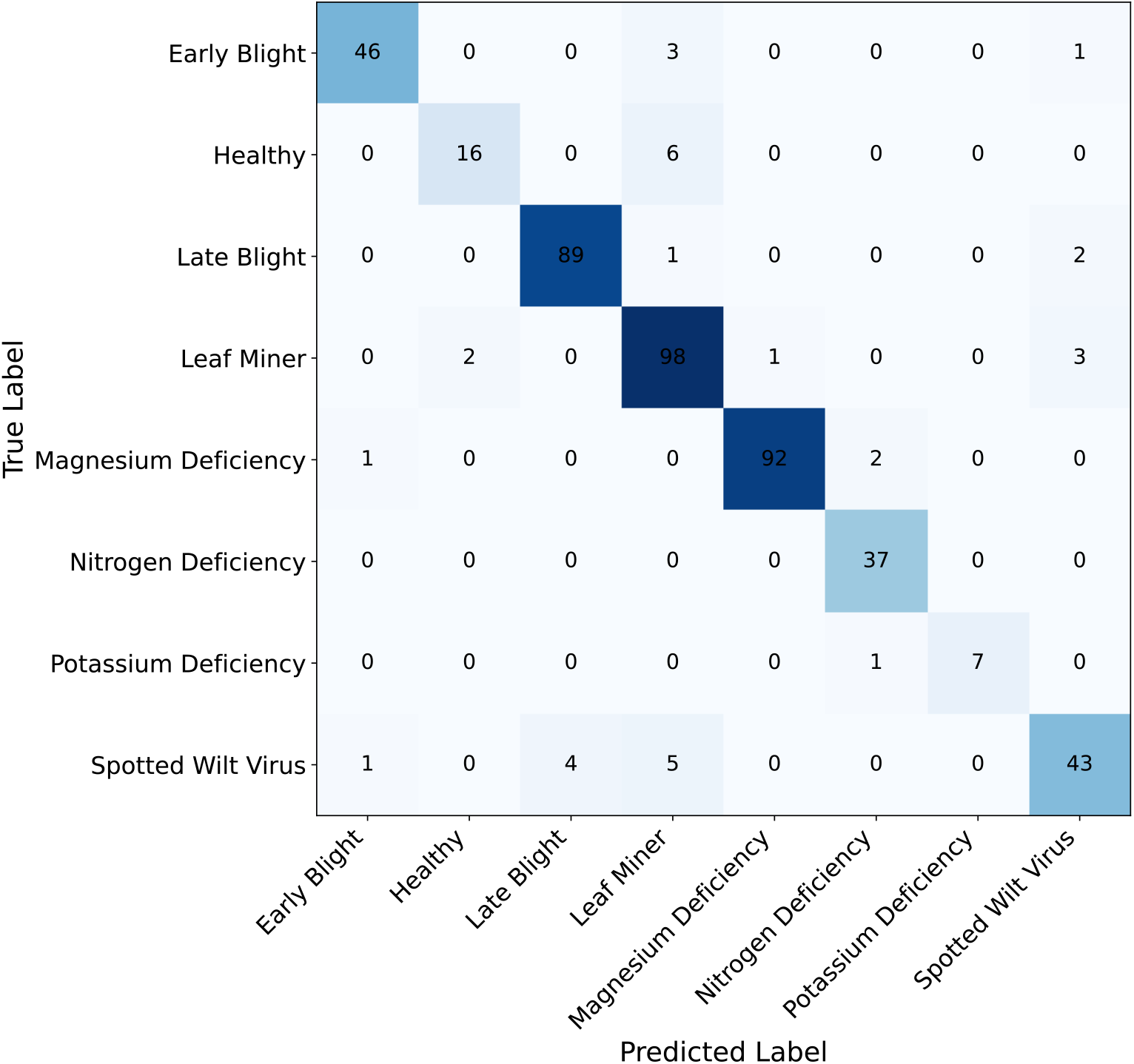
Confusion matrix of the proposed LeafyVGG-16 model for tomato leaf disease and nutrient deficiency classification.

Furthermore, the training and validation accuracy/loss curves shown in Fig. 5 indicate stable convergence behavior throughout the training process. The validation accuracy consistently improved while the validation loss remained comparatively low, suggesting effective feature learning and reduced overfitting capability of the proposed framework.

### B. Comparative Performance Analysis

To further evaluate the effectiveness of the proposed framework, the performance of LeafyVGG-16 was compared with several deep learning architectures, including InceptionV3, DenseNet-121, EfficientNetB0, ResNet-50, and a baseline CNN model. All transfer learning-based models utilized pre-trained ImageNet weights, whereas the CNN model was trained from scratch.

Table II summarizes the comparative performance results in terms of accuracy, precision, recall, and F1-score. Among all evaluated models, the proposed LeafyVGG-16 framework achieved the highest overall classification accuracy of 0.93 and the highest F1-score of 0.92, demonstrating superior classification capability for tomato leaf disease and nutrient deficiency recognition.

**TABLE II.** Comparative performance analysis of different deep learning models.

| Model | Accuracy | Precision | Recall | F1-score |
| --- | --- | --- | --- | --- |
| InceptionV3 | 0.90 | 0.89 | 0.85 | 0.87 |
| CNN | 0.90 | 0.88 | 0.89 | 0.89 |
| DenseNet-121 | 0.92 | 0.91 | 0.89 | 0.90 |
| EfficientNetB0 | 0.78 | 0.74 | 0.72 | 0.71 |
| ResNet-50 | 0.90 | 0.93 | 0.84 | 0.86 |
| **Proposed LeafyVGG-16** | **0.93** | **0.93** | **0.90** | **0.92** |

The proposed model also achieved the highest precision score of 0.93, indicating strong discriminative ability and reduced false-positive predictions. Although ResNet-50 obtained a similar precision value, its recall and F1-score were comparatively lower, suggesting reduced consistency in identifying all disease categories. DenseNet-121 demonstrated competitive performance with an accuracy of 0.92 and F1-score of 0.90, while InceptionV3 and the baseline CNN model achieved overall accuracies of 0.90.

EfficientNetB0 showed the lowest overall performance among the evaluated architectures, achieving an accuracy of 0.78 and F1-score of 0.71. The comparatively lower performance may be attributed to insufficient feature adaptation to the dataset characteristics under the selected training configuration.

Overall, the experimental results indicate that the proposed LeafyVGG-16 framework provides a balanced combination of feature extraction capability, generalization performance, and classification robustness, outperforming the compared deep learning models across most evaluation metrics.

### C. Adversarial Robustness Analysis

To evaluate robustness, adversarial attacks were performed using FGSM and PGD methods. FGSM generates single-step gradient-based perturbations, while PGD applies iterative updates to generate adversarial examples within a defined perturbation constraint.

The performance degradation under FGSM attack is illustrated in Fig. 7, where accuracy gradually decreases from 0.92 to 0.86 as epsilon increases from 0.01 to 0.05.

**Fig. 7.**
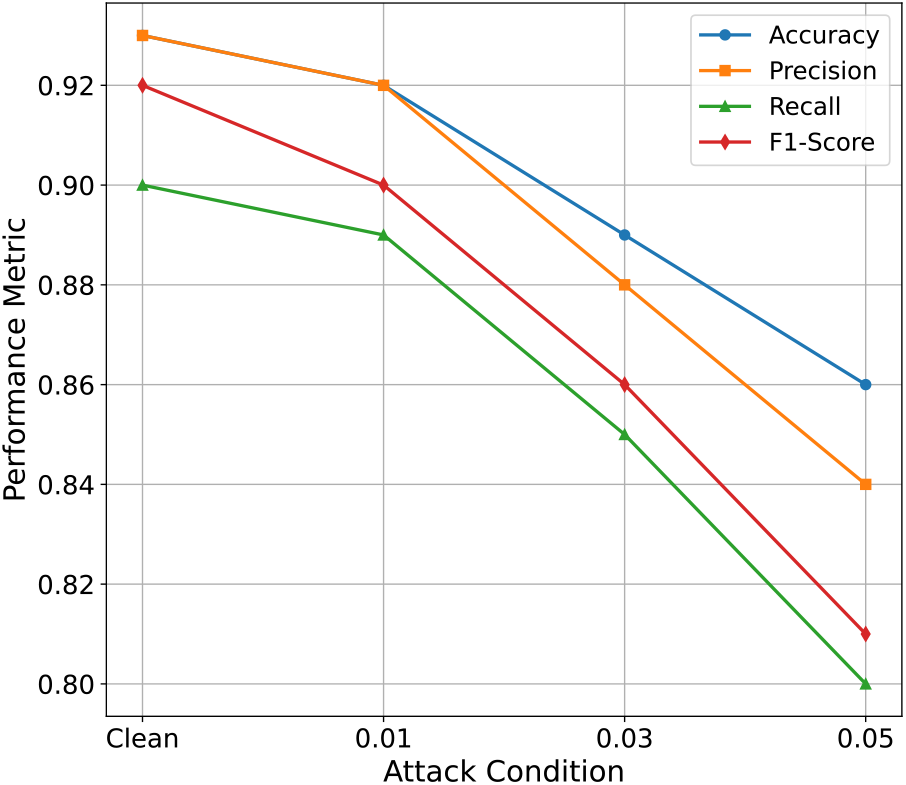
Performance degradation of the proposed LeafyVGG-16 model under FGSM adversarial attacks with increasing perturbation strength.

Under PGD attack (epsilon = 0.05, step size = 0.005, 10 iterations), the model achieves an accuracy of 0.86, precision of 0.84, recall of 0.80, and F1-score of 0.81.

Overall, both FGSM and PGD attacks lead to a noticeable reduction in model performance, indicating that the proposed framework is sensitive to adversarial perturbations. This highlights the importance of improving robustness for real-world agricultural applications where reliability and security are critical.

### D. Discussion

The experimental results demonstrate that the proposed LeafyVGG-16 model achieves strong classification performance compared to several state-of-the-art deep learning architectures. However, the adversarial robustness analysis reveals an important limitation: the model is sensitive to small perturbations in input data.

This observation is critical in the context of agricultural decision-support systems, where deep learning models may be integrated into UAV-based monitoring systems, mobile diagnostic tools, or automated farm management platforms. In such scenarios, adversarial perturbations—whether caused by environmental noise, sensor artifacts, or intentional attacks—can lead to incorrect disease predictions. This may result in improper pesticide application, missed disease detection, or unnecessary agricultural interventions.

The results further indicate that both FGSM and PGD cyber attacks negatively affect the classification capability of the proposed model, suggesting that the learned feature representations are not fully robust to adversarial perturbations. This observation is consistent with existing adversarial machine learning studies, where standard CNN-based architectures without adversarial defense mechanisms show limited resistance to gradient-based attacks.

From a broader perspective, this study highlights an important gap in current agricultural AI systems: most existing research focuses on improving classification accuracy under clean and controlled conditions, while robustness and security considerations remain underexplored. The findings of this work emphasize that high accuracy alone is insufficient for real-world deployment in safety-critical agricultural environments.

## V. Conclusion

This study presented a transfer learning-based framework, LeafyVGG-16, for plant disease and nutrient deficiency classification using real-world tomato leaf images. The proposed model utilizes a deep convolutional architecture enhanced with a customized classification head and is trained using a structured fine-tuning strategy to improve feature representation and generalization capability.

Experimental results demonstrate that LeafyVGG-16 consistently outperforms several state-of-the-art deep learning models, including ResNet-50, DenseNet-121, InceptionV3, EfficientNetB0, and a baseline CNN. The proposed model achieved the highest classification performance, with an accuracy and precision of 0.93, a recall of 0.90, and an F1-score of 0.92, indicating its effectiveness in distinguishing fine-grained plant disease and nutrient deficiency categories.

In addition to standard performance evaluation, the robustness of the proposed model was systematically analyzed under adversarial cyber attack scenarios using FGSM and PGD methods. The results reveal that although LeafyVGG-16 performs strongly under clean conditions, its performance degrades under adversarial perturbations generated by both FGSM and PGD attacks. This highlights an important limitation in the model’s resilience when exposed to gradient-based adversarial attacks, which may pose security and reliability risks in real-world agricultural deployment systems.

Overall, the study demonstrates that while deep transfer learning models such as LeafyVGG-16 are highly effective for plant disease classification, their vulnerability under adversarial conditions must be carefully considered before deployment in practical environments.

Future work will focus on developing cyber-resilient deep learning models for agricultural applications. This includes integrating adversarial training strategies, robust optimization techniques, and defense mechanisms to improve resistance against adversarial attacks. Furthermore, extending the framework to multi-crop datasets and real-time field deployment scenarios will be explored to enhance the reliability and security of AI-driven precision agriculture systems.

## Acknowledgment

This work was supported by the Evans-Allen Capacity Grant from the U.S. Department of Agriculture National Institute of Food and Agriculture. This work was also supported by the West Virginia Higher Education Policy Commission (HEPC), Science, Technology and Research (STaR) Division under Grant No. IGP25-001.

